# Comparative genomics reveals shared accessory regions between members of two *Fusarium* species complexes virulent on garden pea

**DOI:** 10.64898/2026.06.29.735274

**Authors:** Ambika Pokhrel, Sajeet Haridas, Sara Calhoun, Alan Kuo, Anna Lipzen, Robert Riley, Kurt LaButti, Jasmyn Pangilinan, Bill Andreopoulos, Guifen He, Mi Yan, Kerrie Barry, Li-Jun Ma, David M. Geiser, Michael Freitag, Igor V. Grigoriev, Jeffrey J. Coleman

## Abstract

The contribution of accessory or conditionally dispensable chromosomes to host-specific virulence was first demonstrated in members of the *Fusarium solani* species complex (FSSC) that are pathogens of garden pea, *Pisum sativum* L. The phenomenon has since been shown to exist in many fungal plant pathogens, including the closely related *F. oxysporum* species complex (FOSC). Genome analysis of members of the FSSC and FOSC pathogenic on pea revealed a diverse size range of the accessory genome of these fungi. Despite the ∼65 million years of diverging time, regions on a chromosome known to carry host-specific virulence factors for pea, including the cytochrome P450 pisatin demethylase (*PDA*) and other pea pathogenicity (*PEP*) genes, were present in all genomes of these pea pathogens. Genes directly involved in virulence on pea – *PEP2*, *PDA*, and *PEP5*– were the most frequently clustered together. Transcriptome analysis of fungal mycelia treated with the pea phytoalexin pisatin, identified 1,155 differentially expressed genes where many were involved in cellular stress responses. As wilt pathogens that invade host xylem, members of the FOSC encode more putative effectors, when compared to those in the FSSC, and several FOSC effectors were identified to confer race specificity. The conservation of part of the accessory genomes across two evolutionarily diverged species complexes suggests a common origin. Horizontal transfer of accessory chromosomes containing genetic loci involved in pathogenesis for garden pea offers a parsimonious explanation of the polyphyletic origin of host specificity.

## Introduction

Members of the *Fusarium solani* and *F. oxysporum* species complexes (FSSC and FOSC, respectively) comprise soil-borne filamentous fungi of agricultural importance (Coleman 2016; Michielse and Rep 2009; Geiser et al. 2021). Collectively, members of the FSSC and FOSC have a broad host range capable of infecting hundreds of cultivated plant hosts, where *Fusarium vanettenii* (*Fv*; previously known as *Nectria haematococca* MP VI and *Fusarium solani* f. sp. *pisi*) and *Fusarium oxysporum* f. sp. *pisi* (*Fop*), members of the FSSC and FOSC, respectively, cause *Fusarium* root rot and wilt diseases on garden pea (*Pisum sativum* L.).

Phylogenomic studies revealed *Fusarium* to be monophyletic (Geiser et al. 2021), however, host specificity in *Fusarium* is a polyphyletic trait (Van Dam et al. 2018). The polyphyletic origin of host specificity is at least partially due to the presence of genes that reside in dispensable regions or entire chromosomes which are capable of being laterally transferred between isolates of *Fusarium* (Miao et al. 1991; Coleman et al. 2009; Ma et al. 2010; Van Dam et al. 2018). Thus, the genome of members of the FSSC and FOSC can be divided into “core” and “accessory” regions where the core genome is conserved and usually syntenic between a majority of the isolates and harbors essential genes for the fungus, while the accessory genome is unique and diverse (Coleman et al. 2009; Ma et al. 2010). Core genes are generally shared widely across broad taxonomic categories and show evolutionary patterns consistent with organismal history, while accessory genes tend to have sporadic distributions, even within genera and species. Accessory genes and chromosomes (also known as supernumerary, lineage-specific, and in some instances conditionally dispensable chromosomes) are important for fungal adaptation and pathogenicity because they often harbor various virulence related genes as well as traits important for fungal adaptation and niche colonization (Miao et al. 1991; Rodriguez-Carres et al. 2008; Schmidt et al. 2013).

The first studies demonstrating the contribution of the fungal accessory genome to host-specific virulence were with *Fv* (Miao et al. 1991). The ability of the fungus to degrade the pea phytoalexin (+)pisatin is conferred by the cytochrome P450 pisatin demethylase (Pda) encoded by a gene located on an accessory chromosome. This accessory chromosome has been described as a conditionally dispensable (CD) chromosome, as the 1.6 Mb chromosome is not essential for the fungus, although when present conferred host specific virulence towards garden pea. Other genes involved in conferring host specificity towards garden pea also reside on the chromosome, referred to as the *PDA1* chromosome (Han et al. 2001; Temporini and VanEtten 2002, 2004). These four additional pea pathogenicity (*PEP*) genes, *PEP1, PEP2, PRG1* (*cDNA3*), and *PEP5*, are organized in a ∼20 kb locus with *PDA1* and is known as the *PEP* cluster (Han et al. 2001; Pokhrel and Coleman 2024). Besides the *PEP* genes contributing to pea pathogenicity, the *PDA1* chromosome is also known to harbor homoserine utilization (*HUT*) genes that provide a competitive advantage for colonization of the pea rhizosphere (Rodriguez-Carres et al. 2008). The first sequenced genome of *Fv* (strain 77-13-4), revealed at least three chromosomes (14, 15, and 17) were accessory (Coleman et al. 2009). Outside of *Fv*, homologs of *PDA* and the *PEP* genes have been identified in *Fop* as well as *Fusarium tenuicristatum*, another member of the FSSC (Temporini and VanEtten 2004; Coleman et al. 2011).

Seeking to understand the diversity, as well as conservation of the accessory genomes and *PDA* chromosomes in *Fv* and *Fop* field isolates, the genomes of sixteen isolates of *Fv,* five isolates of *Fop,* and an isolate of *F. tenuicristatum* collected from various plant hosts and geographic locations were sequenced. Fungal accessory genomes are known to be dynamic and rapidly evolving, and while there was diversity found among the accessory genomes of these Species Complexes, there were shared regions - in particular those associated with the accessory chromosome that harbors *PDA*. While this partly explains virulence on the common host, garden pea, it also suggests a shared ancestral origin of these accessory regions independent of the common ancestry of the FSSC and FOSC, which may have diverged as long as ∼65 million years ago (O’Donnell et al. 2013).

## Results

### Phylogeny and genome features of FSSC and *Fop* isolates

The phylogenetic relationship between a collection of FSSC and *Fop* isolates (Supplemental Table S1) was evaluated using the entire coding sequence of three genes, *TEF1*, *RPB1*, and *RPB2*. These three gene sequences were concatenated and a maximum-likelihood phylogenetic tree was constructed where clade support was assessed by 1,000 bootstrap replicates (Fig. 1A). In the phylogenetic analysis, the isolates were divided into two clades corresponding to each species complex. The *Fv* isolates grouped together with other members of the FSSC (*F. mori* NRRL 22230 and *F. tenuicristatum* NRRL 22470) and the *Fop* isolates grouped separately (Fig. 1A).

**Fig. 1:**
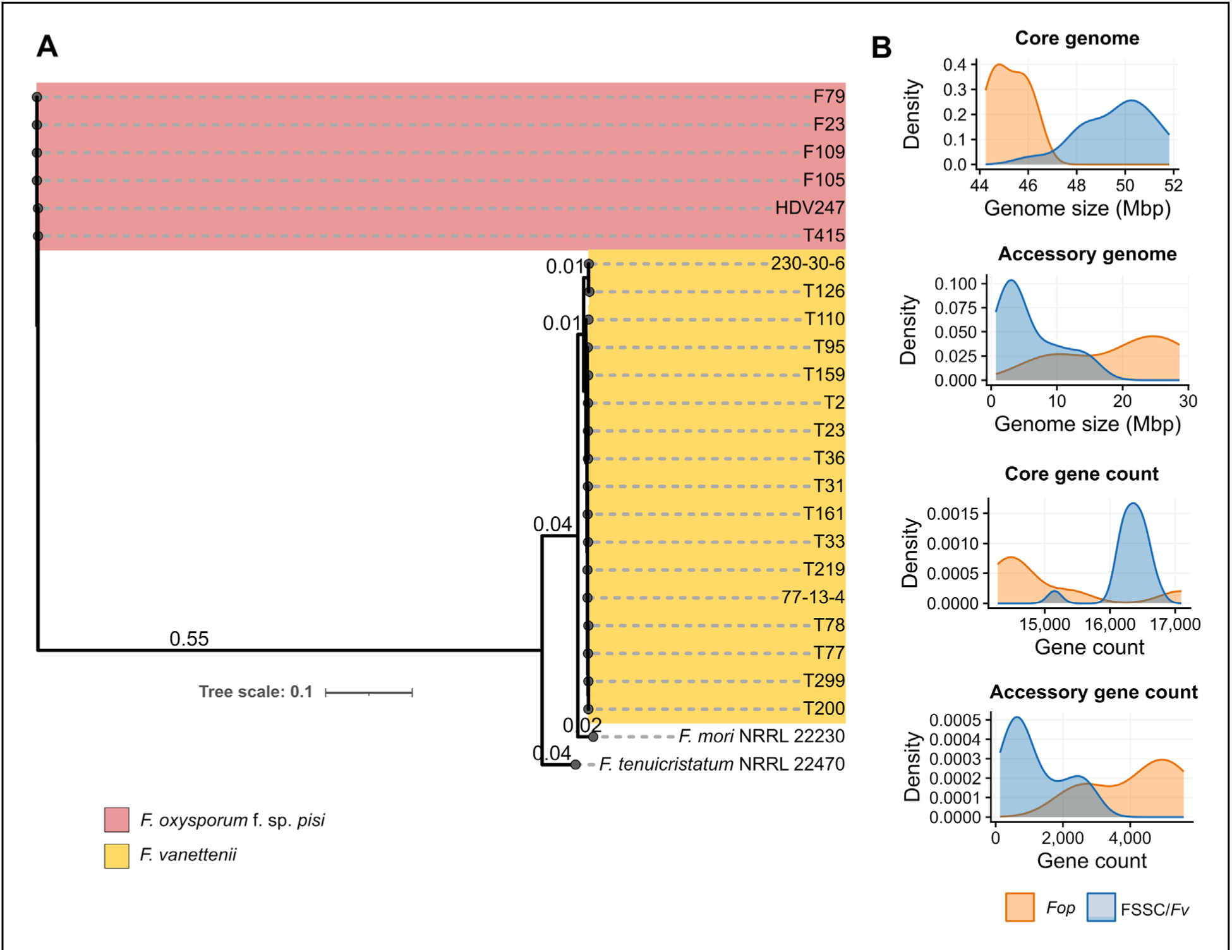
Phylogeny and basic genome features of FSSC and *Fop* isolates. **(A)** Maximum-likelihood phylogenetic tree of FSSC and *Fop* isolates constructed using *TEF1*, *RPB1*, and *RPB2* gene sequences. Branch support values are shown at corresponding nodes. The clade in pink represents isolates of *F. oxysporum* f. sp. *pisi* and the clade in yellow represents isolates of *F. vanettenii*. **(B)** Histograms showing the distribution of genome size (core and accessory) and gene count (core and accessory) in FSSC and *Fop* isolates.

The size of the seventeen *Fv* genomes included in this study ranged between that of isolate T110 (non-pathogenic to pea; 48.93 Mb) and isolate T36 (63.21 Mb) (Table1; Supplemental Table S2), while the genome size of the pea pathogenic *F. tenuicristatum* isolate NRRL 22470 was 61.51 Mb, larger than most of the genomes of the *Fv* isolates. The average genome size of the *Fop* isolates was greater than that of the FSSC isolates (64.98 Mb versus 55.45 Mb), with 55.19 Mb being the smallest (HDV247) and 73.3 Mb the largest (F23) (Table 1; Supplemental Table S2). The GC content of all isolates was ∼48 to 50% (Table1; Supplemental Table S2). The total number of genes in the FSSC genomes ranged from approximately 16,000 to 18,000 with the least in isolate T110 (16,335). The average number of genes in the *Fop* isolates was greater than in the FSSC isolates with the highest number in isolate F23 with 20,063 and lowest in isolate T415 with 18,001 (Table 1; Supplemental Table S2).

**Table 1.** Genome features of the core and accessory genome of FSSC and *Fop* isolates used in this study.

| Species | Isolate | Genome size (Mbp) |  |  | GC content (%) |  |  | Number of genes |  |  |
| --- | --- | --- | --- | --- | --- | --- | --- | --- | --- | --- |
|  |  | Total | Core | Accessory | Total | Core | Accessory | Total | Core | Accessory |
| FSSC | T2 | 60.99 | 51.47 | 9.52 | 48.75 | 50.79 | 45.99 | 18,837 | 16,562 | 2,275 |
|  | T23 | 61.89 | 48.01 | 13.88 | 49.45 | 49.77 | 46.68 | 18,783 | 16,086 | 2,697 |
|  | T31 | 52.05 | 50.95 | 1.11 | 49.21 | 49.46 | 37.83 | 16,729 | 16,503 | 226 |
|  | T33 | 54.72 | 50.36 | 4.36 | 49.53 | 49.95 | 44.82 | 17,498 | 16,586 | 912 |
|  | T36 | 63.21 | 50.3 | 12.91 | 48.92 | 49.34 | 47.34 | 18,785 | 16,155 | 2,630 |
|  | T77 | 54.39 | 50.56 | 3.83 | 49.38 | 49.63 | 46.26 | 17,174 | 16,265 | 909 |
|  | T78 | 58 | 49.19 | 8.82 | 49.6 | 50.1 | 46.84 | 18,014 | 16,212 | 1,802 |
|  | T95 | 51.64 | 48.8 | 2.84 | 49.42 | 50.16 | 36.89 | 16,846 | 16,421 | 425 |
|  | T110 | 48.93 | 48.22 | 0.71 | 49.47 | 49.69 | 35.31 | 16,335 | 16,205 | 130 |
|  | T126 | 57.33 | 48.44 | 8.89 | 49.31 | 50.14 | 44.82 | 18,069 | 16,359 | 1710 |
|  | T159 | 53.28 | 49.97 | 3.31 | 48.78 | 49.6 | 36.45 | 17,252 | 16,736 | 516 |
|  | T161 | 53.46 | 51.39 | 2.07 | 49.37 | 49.42 | 48.27 | 17,043 | 16,466 | 577 |
|  | T200 | 52.49 | 48.08 | 4.41 | 50.34 | 50.04 | 45.53 | 17,115 | 16,315 | 800 |
|  | T219 | 52.6 | 49.97 | 2.63 | 49.8 | 50.25 | 45.22 | 17,132 | 16,575 | 557 |
|  | T299 | 54.43 | 50.2 | 4.22 | 49.55 | 49.26 | 46.98 | 17,256 | 16,206 | 1,050 |
|  | 77-13-4 | 54.59 | 51.82 | 2.77 | 48.91 | 49.25 | 42.61 | 16,929 | 16,370 | 559 |
|  | 230-30-6 | 52.62 | 49.9 | 2.72 | 49.15 | 49.57 | 45.21 | 17,092 | 16,402 | 690 |
|  | NRRL 22470 | 61.51 | 46.18 | 15.33 | 47.28 | 50.8 | 36.64 | 17,800 | 15,151 | 2,649 |
| *Fop* | F23 | 73.3 | 44.66 | 28.65 | 48.24 | 48.45 | 47.92 | 20,063 | 14,471 | 5,592 |
|  | F79 | 67.02 | 44.24 | 22.77 | 48.35 | 48.52 | 48.06 | 19,180 | 14,554 | 4,626 |
|  | F105 | 71.47 | 45.49 | 25.97 | 48.3 | 48.4 | 48.14 | 19,358 | 14,278 | 5,080 |
|  | F109 | 67.09 | 44.86 | 22.23 | 48.24 | 48.35 | 48.04 | 19,272 | 14,580 | 4,692 |
|  | HDV247 | 55.19 | 45.87 | 9.31 | 47.04 | 47.71 | 47.11 | 19,623 | 17,094 | 2,529 |
|  | T415 | 55.84 | 46.28 | 9.56 | 46.39 | 47.66 | 46.84 | 18,001 | 15,388 | 2,613 |

### Core and accessory genomic regions in the *Fop* and FSSC isolates

Comparison of the genomes to well referenced *Fusarium* genomes enabled identification of core and accessory scaffolds. The majority (∼60 - 98.5%) of the *Fop* and FSSC genomes was identified as the core genome (Table 1; Supplemental Fig. S1), with *Fv* isolate T110 possessing only 707,300 bp (1.44% of the total genome) predicted to be accessory (Table 1; Supplemental Fig. S1). The GC content was lower in the accessory genomes when compared to the corresponding core genomes, as low as 35.3% (T110) following a trend previously observed (Coleman et al., 2009); in contrast in *Fop* genomes the lowest accessory genome GC content was 46.8% (T415) (Table 1; Supplemental Fig. S1). The total number of core genes in the FSSC genomes was ∼16,000, except for the *F. tenuicristatum* NRRL 22470 genome, which was predicted to encode 15,151 core genes. However, the number of predicted genes in the accessory genome was highly variable, ranging from as little as 130 (T110) to 2,697 (T23) (Table 1; Supplemental Fig. S1-B). Among the *Fop* genomes, the highest number of core genes was found in HDV247 (17,094), while the highest number of accessory genes was in F23 (5,592) (Table 1; Supplemental Fig. S1A). Overall, while the core genome size in FSSC isolates was larger than in *Fop* genomes, the accessory genome size was significantly higher in *Fop* isolates (Fig. 1B). This structural divergence corresponded to the gene content; the number of core genes was consistently high in FSSC isolates compared to *Fop* isolates, whereas the number of predicted accessory genes were markedly higher in *Fop* genomes (Fig. 1B).

GO enrichment analysis of the core and accessory proteins from the FSSC and *Fop* isolates was performed by comparison against the *Fv* 77-13-4 v2 (Coleman et al. 2009) and the *F. oxysporum* f. sp. *lycopersici* 4287 proteomes (Ma et al. 2010; Ayhan et al. 2018), respectively. The significantly enriched GO pathways were classified into three main categories: biological process (BP), molecular function (MF), and cellular component (CC) (Fig. 2; Supplemental Fig. S2). Overall, the GO enrichment results indicate the core proteins play an important role in the growth and development of fungi. This is evident where > 100 GO pathways from the BP and CC categories including those in biosynthetic and metabolic processes, maintaining structure of various cellular and intracellular organelles, cytoplasm, and protein containing complexes were represented by core proteins in the *Fop* and FSSC genomes. On the contrary, proteins from GO pathways in the MF category were abundant in the accessory genomes representing enzymes with lyase, antioxidant, hydrolase, and various ion binding activities. These activities that are not necessarily required for survival, growth, and development of fungi, but undertaken when necessary. This is further evident with the scarcity of significantly enriched CC pathways for the accessory genomes of both *Fop* and FSSC isolates, supporting the notion accessory proteins may not be intricately involved in functions related to cellular structure, location, or organelle development (Navasca et al. 2025).

**Fig. 2:**
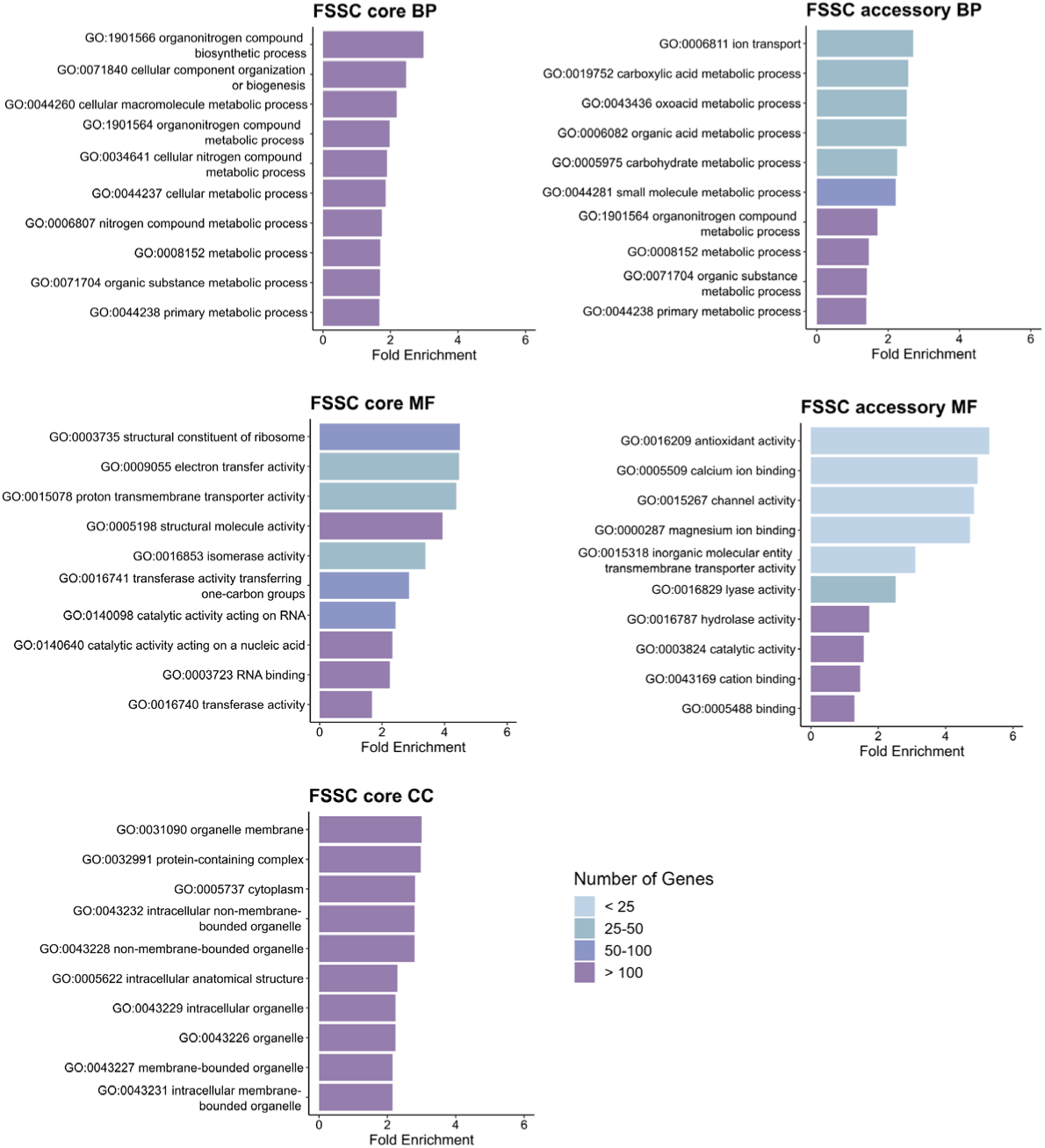
Gene enrichment analysis of core and accessory genomic regions of FSSC isolates. Top ten significantly enriched gene ontology pathways of the core and accessory genome in FSSC isolates. The statistical significance of gene enrichment was determined at FDR threshold < 0.05. The enriched pathways are categorized as biological process (BP), molecular function (MF), and cellular components (CC). *Only one gene ontology pathway under cellular component category ‘extracellular region’ was significantly enriched in the FSSC isolates.

The transposable element (TE) content in the *Fop* and FSSC genomes directly correlated with the genome size (Fig. 3A; Supplemental Table S3). TE content in the *Fop* genomes averaged 11.11% and ranged from 3.91% (T415) to 15.56% (F105); whereas the TE content in the FSSC genomes ranged from 5.34% (T110) to 14.09% (NRRL 22470) (Supplemental Table S3), with an average of 7.76% of the genome. The elevated TE content in NRRL 22470 when compared to the other FSSC genomes was primarily due to the greater incidence of class I (LTR- Unknown), class II transposons (PIF-Harbinger and Mutator), and ‘unknown’ repetitive elements (Supplemental Table S3; Fig. 3C). Overall, the TEs predicted in the *Fop* and FSSC genomes showed a significant number of ‘unknown’ repetitive elements. In *Fop* genomes the TE content in the core genome ranged from 2.61% (HDV247) to 6.62% (F109) of the core genome, whereas in the accessory genome it ranged from 14.57% (T415) to 35.83% (F23) of the accessory genome (Fig. 3B). The majority of the TEs in accessory genome of F23 (13.56%) were classified as ‘unknown’ repetitive elements. In FSSC genomes the TE content in the core genome ranged from 2.98% (T126) to 7.51% (T2) of the core genome, whereas in accessory genome it ranged from 1.07% (T110) to 37.57% (NRRL22470) of the accessory genome (Fig. 3C). T110 is comprised of only a single class of TEs in the accessory genome, class II (Helitron). The enrichment of TEs in the accessory genomic regions may contribute to genome plasticity and acquisition of virulence associated genes or loci as reported in other *Fusarium* species (López Díaz et al. 2025; Yang et al. 2020).

**Fig. 3:**
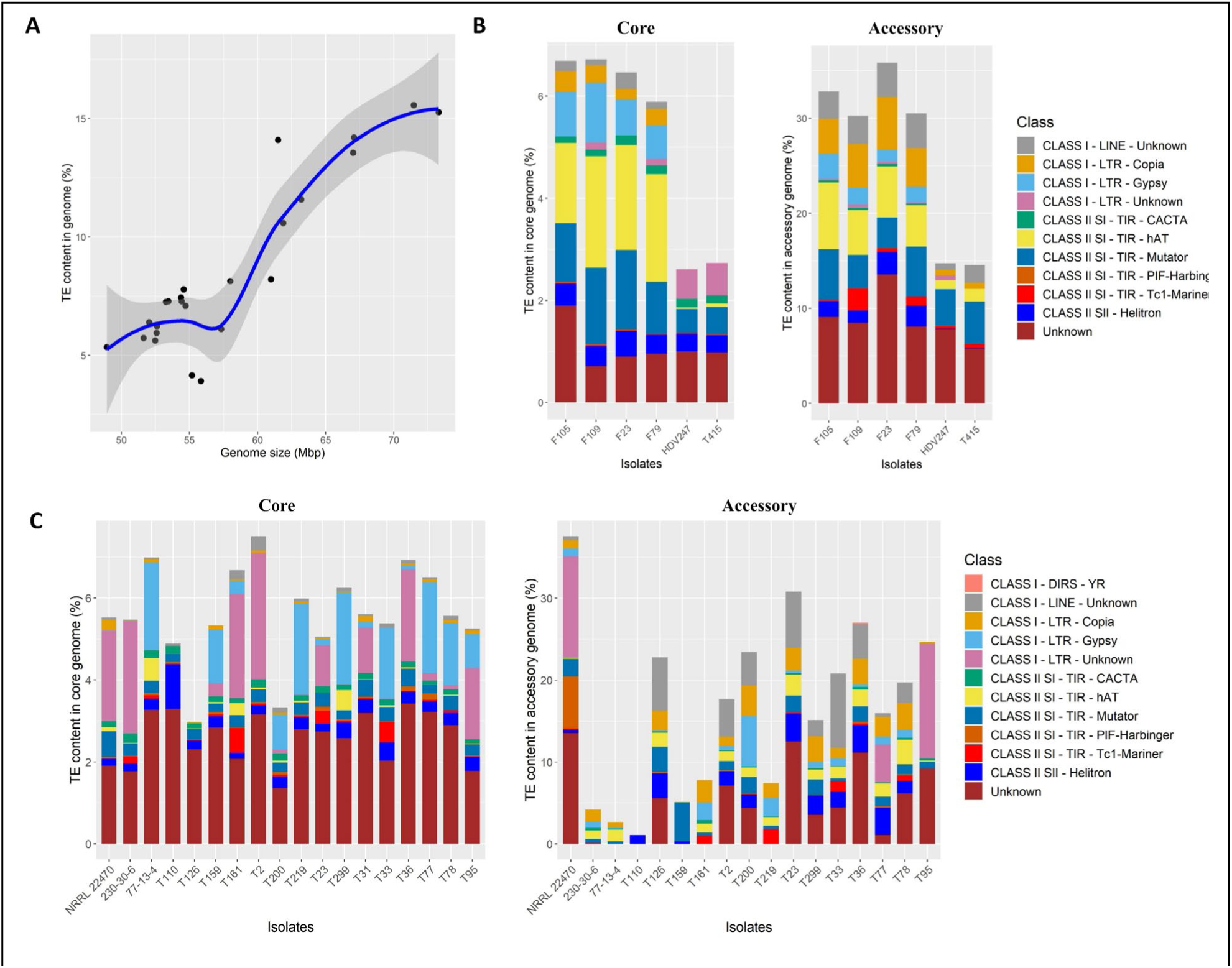
Transposable element (TE) content and genome size variation in *Fop* and FSSC isolates. **(A)** Contribution of TEs in total genome size across 24 *Fop* and FSSC isolates. **(B)** Contribution of each TE class to genome size for *Fop* isolates. **(C)** Contribution of each TE class to genome size for FSSC isolates.

### Conservation of accessory genomes between the FSSC and FOSC pea pathogens

The accessory genomes of the FSSC and *Fop* isolates were compared to the established reference accessory genome of *Fv* 77-13-4. Conservation of the *PDA1* encoding accessory chromosome 14 was evident between the accessory genomes of the pea pathogenic FSSC isolates (Fig. 4A), where this conservation was higher between the FSSC genomes when compared to the accessory genome of *Fop* (Fig. 4B). Specifically, the largest contiguous conserved block on chromosome 14 in the FSSC spanned ∼58.4 kb with 97.76% identity (observed in isolates such as T200 and T299), which is nearly seven-fold longer than the maximum conservation observed in the *Fop* isolates (∼8.0 kb with 93% identity) (Fig. 4B). Furthermore, examination of other accessory chromosomes, chromosome 16 (scaffold_16) and chromosome 17 (scaffold_17) showed conservation in the FSSC and *Fop* genomes. Within the FSSC, the largest conserved region on chromosome 16 was ∼28.9 kb (98.96% identity; with isolate T159), and on chromosome 17 it was ∼48.8 kb (98.16% identity; with isolate T77). In stark contrast, alignments to these accessory chromosomes in the *Fop* genomes yielded significantly smaller fragmented regions. The maximum contiguous conservation in *Fop* was restricted to blocks of only ∼2.2 kb (73.52% identity) for chromosome 16 and ∼4.3 kb (65.55% identity) for chromosome 17 (Fig. 4B).

**Fig. 4:**
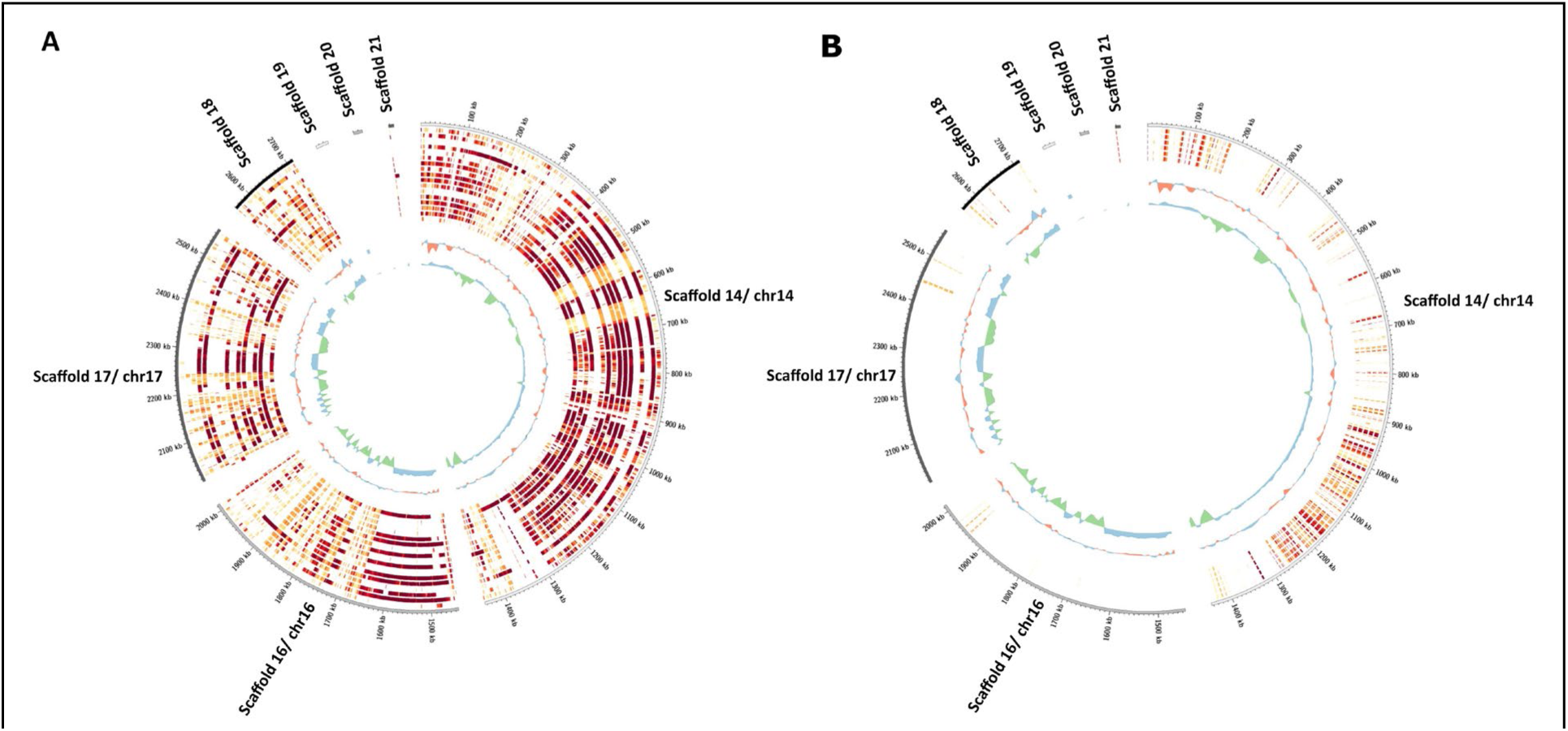
Conservation of accessory genomes between FSSC and FOSC. **(A)** Circos heatmap showing conservation of the accessory genome between FSSC isolates, where the accessory genome from *Fv* 77-13-4 was used as a reference for comparison. Datapoints with more than 75% similarity are shown, and light yellow indicates less similarity and red indicates highly similar regions. The order of genomes starting from outer ring is 77-13-4, NRRL 22470, 230-30-6, T110, T126, T159, T161, T200, T219, T23, T299, T2, T31, T33, T36, T77, T78, and T95. Green and blue lines indicate GC% variance (value< 0 = green; value> 0 = blue). Red and blue lines indicate GC% skew (value< 0 = red; value> 0 = blue). **(B)** Circos heatmap showing conservation of accessory scaffolds between *Fop* genomes and *Fv* 77-13-4. Datapoints with more than 75% similarity are shown, and light yellow indicates less similarity and red indicates highly similar regions. The order of genomes starting from outer ring is 77-13-4, F105, F109, F23, F79, HDV247, and T415.

Even though *Fv* and *Fop* isolates belong to different species complexes (Fig. 1A), there is some conservation of the accessory genome between these two groups, specifically for chromosome 14 which is linked to host-specificity on pea. Further analysis of these conserved regions between *Fop* and FSSC genomes identified that the largest region shared was ∼10 kb between *Fop* genomes (F23, F79, and F105) and FSSC genomes (T2, T23, and T36) (Supplemental Table S4). Notably, this region was composed of a single annotated gene categorized as a ‘hypothetical protein’ and exhibits high sequence identity (∼99%) across these distinct genomes (Supplemental Data S1). Despite its conservation, the region is gene-poor and structurally repetitive, suggesting it may represent a TE-associated segment or a low-complexity region. These findings suggest that while some accessory genomic regions appear conserved between FSSC and *Fop* isolates, they may be largely repetitive in nature, potentially highlighting the role of repetitive elements or TEs in shaping accessory genome architecture and facilitating the acquisition or translocation of virulence-related genes or loci.

### Diversity in *PDA* scaffolds of FSSC isolates

Except for T110 which lacks *PDA*, the FSSC *PDA* containing scaffolds were identified (Supplemental Table S5) and compared to each other to identify syntenic regions (Supplemental Fig. S3A), revealing four groups with significant syntenic regions. The first group contained isolates 230-30-6, T77, T78, T161, T200, T219, and T299 (Supplemental Fig. S3B), with syntenic loci of up to 1.29 Mb in length. Within this group, the *PDA* encoding scaffolds of isolates 230-30-6, T77, and T78 were well conserved with a single inversion in the *PDA* scaffold of T77 (Supplemental Fig. S3B). Specifically, isolate 230-30-6 and T77 shared 423.74 kb with synteny, 230-30-6 and T78 shared 396.97 kb synteny, and T77 and T78 shared 447.27 kb of synteny between them. Similarly, a large syntenic block was shared between isolates T161 and T299 (1.29 Mb, 92.42 % nucleotide identity), and isolates T200 and T219 (350.33 kb, 88.77 % nucleotide identity) (Supplemental Fig. S3B). Three other groups had synteny between the *PDA* scaffolds: T23 and T36 (213.04 kb), T33 and T126 (298.04 kb), and T31 and T159 (1.41 Mb) (Supplemental Fig. S3C-E). Interestingly, the *PDA* scaffolds in the fourth group of T31 and T159 were ‘scaffold_6’ and ‘scaffold_4’, respectively, which were identified as part of the core genome. This suggests that a translocation event may have occurred, resulting in accessory regions being located in subtelomeric regions of core chromosomes. Other regions with synteny were also identified between accessory *Fv* scaffolds (Supplemental Fig. S4).

### The *PEP* cluster is conserved in FSSC isolates

Analysis of the *PEP* genes was conducted in all FSSC and *Fop* genomes. A homolog of *PDA* was found in all pathogenic FSSC and *Fop* genomes with the exception of T110, which is non-pathogenic on pea (Fig. 5; Supplemental Table S6). Among the *PEP* genes, homologs of *PEP1* were found in all FSSC and *Fop* genomes but were not necessarily found on the scaffold encoding *PDA* (Fig. 5; Supplemental Tables S6 and S7). *PRG1* (*cDNA3*) was only found in *Fv* isolates with the exception of *F. tenuicristatum* NRRL 22470. In isolate T95 (from tulip tree), the *PDA* gene was truncated (782 bp out of 2,088 bp) with 70% nucleotide identity to the functional *PDA1* gene over that region and likely results in the weak virulence on garden pea observed with this isolate (Funnell et al. 2001). Because no other *PEP* genes were present in the same scaffold, it was removed from subsequent analysis. Microsynteny analysis of the *PEP* genes in the *PDA* encoding scaffolds revealed that the gene order is different in FSSC isolates when compared to the originally described *PEP* cluster (Han et al. 2001) (Fig. 5A). The *PEP2*, *PDA*, and *PEP5* genes were the most conserved in the cluster, while the co-occurrence of *PEP1* in the cluster was rare. Even when present, *PEP1* was further removed from the rest of the cluster in the *PDA* scaffold relative to the other *PEP* genes (Fig. 5A). Microsynteny analysis of the *PDA* scaffolds of the *Fop* isolates found that the *PEP* genes were usually not within close proximity of *PDA* in the *Fop* genomes, although homologs of *PEP1* on the *PDA* scaffolds were discovered in isolates HDV247 and F23 (Fig. 5B). Homologs of *PEP1* and *PEP2* were also identified in other non-*PDA* containing scaffolds.

**Fig. 5:**
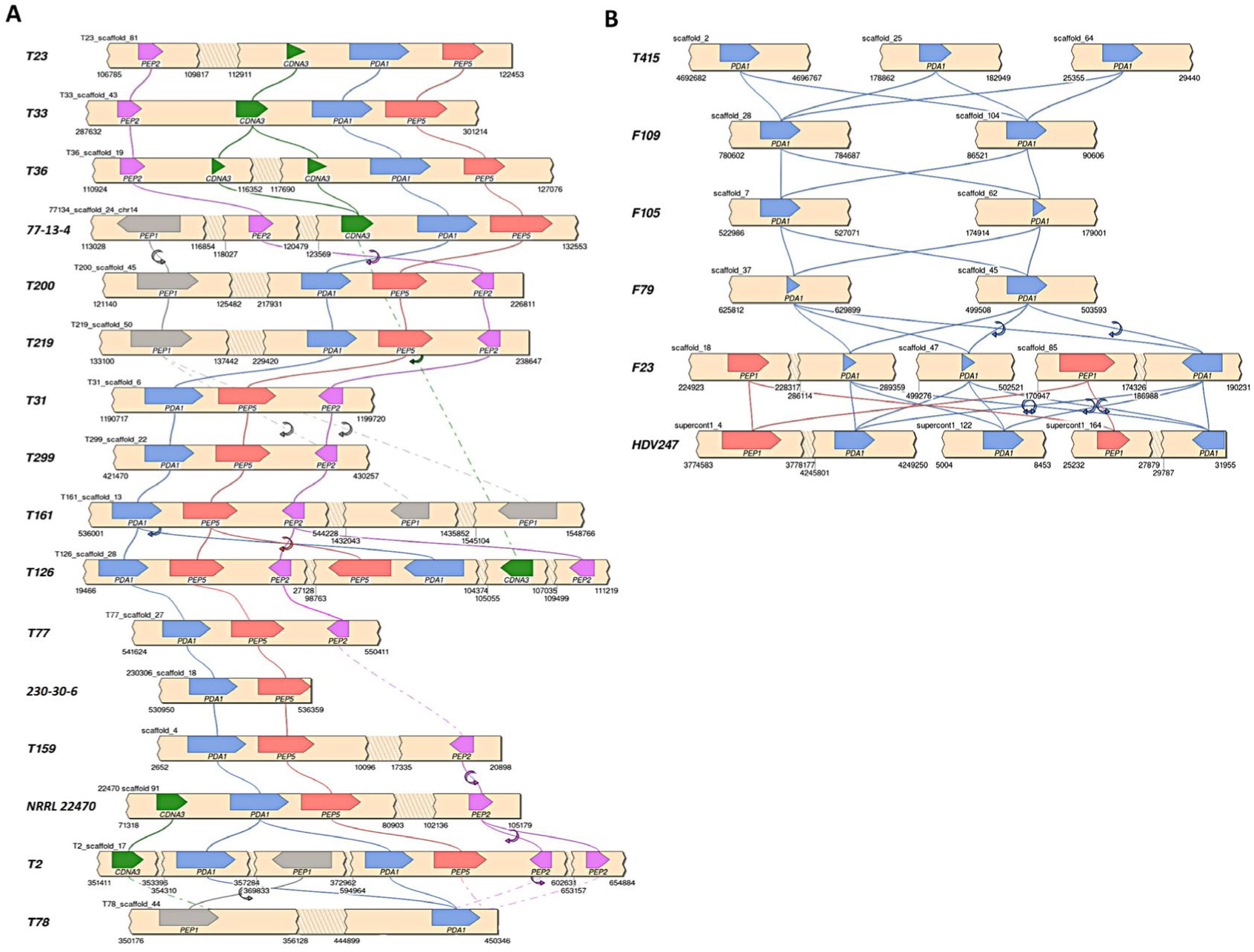
Microsynteny analysis of genes in the *PEP* cluster in the *PDA* scaffolds of (A) FSSC and (B) *Fop* isolates. Each color represents different genes with arrows indicating their orientation in the scaffold. Scaffold names and genomic coordinates are indicated above and below each bar, respectively. Corresponding gene names are indicated below each arrow. Figures were generated using the program SimpleSynteny.

### Identification of host-specific and pathogenicity related effectors and virulence factors in *Fop* and FSSC isolates

The average numbers of predicted effectors in the *Fop* genomes was greater than in the FSSC (Supplemental Table S8). The core genome of the FSSC isolates encoded predicted effectors in the range of 344 (NRRL 22470) to 404 (77-13-4), while the number of putative effectors in the accessory genomes of the isolates was highly variable ranging from 1 (T31 and T110) to 58 (NRRL 22470) (Supplemental Table S8). The number of effectors in the accessory genomes of the *Fop* isolates was relatively higher than in their FSSC counterparts, with the accessory genome of F23 being particularly rich, encoding 29.3% (169) of the total effectors identified (Supplemental Table S8).

Of the total 697 predicted effectors in the *Fop*, 304 were common between all the *Fop* genomes (Fig. 6A). As *Fop* has four defined races (1, 2, 5, and 6), the effectors from the *Fop* genomes were further classified into orthogroups using Orthofinder (Emms and Kelly 2019) in order to identify candidate effector proteins specific to each particular race (Fig. 6A). There were ten conserved putative effectors that were only identified in the race 1 genomes (F79 and F109), 19 in the race 2 genome (T415), and eight in the race 6 genome (F105) (Fig. 6A). There were no putative effectors that were specific and shared between the race 5 genomes (F23 and HDV247), although 38 putative effectors were found exclusively in F23 and 26 exclusively in HDV247. Most of these unique race-specific candidate effectors were uncharacterized with no functional domains, although one of the race 2 specific effectors contained a glycosyl hydrolase 47 family domain and one of the race 6 specific effectors carried a lactonase domain (Supplemental Table S9).

**Fig. 6:**
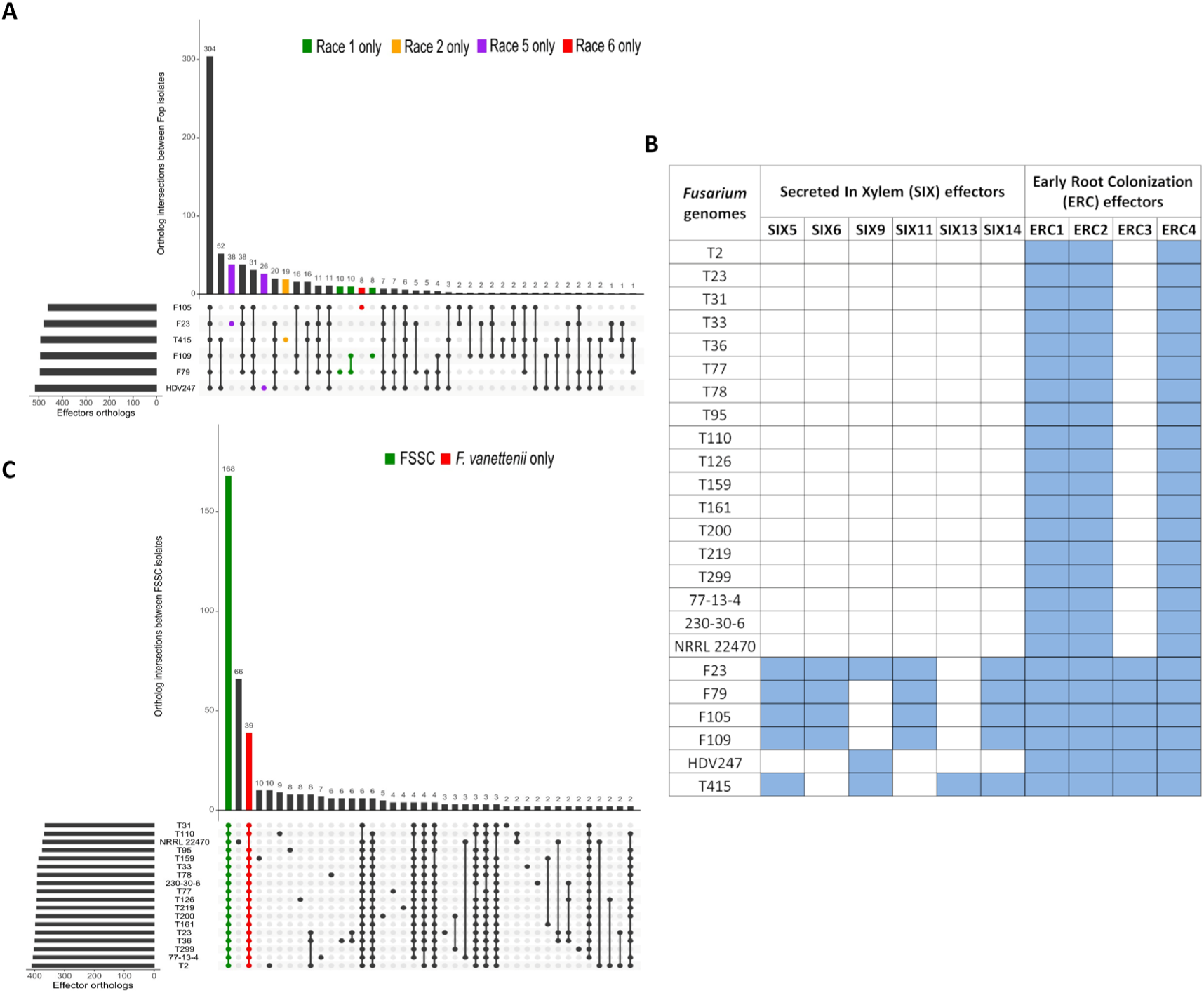
Prediction and distribution of effector proteins in *Fop* and FSSC isolates. **(A)** The number above each bar represents the orthologs of effectors shared between the indicated *Fop* genomes. Bars highlighted in different colors represent race specific groups of effectors in *Fop* genomes that could influence cultivar specificity. **(B)** Chart showing the presence/absence of Six and Erc effectors in *Fop* and FSSC isolates. **(C)** The number above each bar indicates the number of orthologs encoding putative effectors shared between the FSSC isolates. Bars highlighted in different colors are the orthogroups of effectors conserved in various FSSC and *Fv* isolates.

Homologs encoding the well-established secreted in xylem (Six) effector proteins were identified in the *Fop* genomes for *SIX5*, *SIX6*, *SIX9*, *SIX11*, *SIX13*, and *SIX14* (Fig. 6B; Supplemental Table S10); while none were found in the FSSC genomes. Additionally, putative homologs encoding the early root colonization (ERC) effector proteins were found in both FSSC and *Fop*. *ERC1*, *ERC3*, and *ERC4* were identified in both the FSSC and *Fop* genomes, while only *Fop* harbored *ERC2* (Fig. 6B, Supplemental Table S11). Among the ERC effectors, multiple homologs for *ERC1* and *ERC4* were found in the *Fop* and FSSC genomes.

Within all the FSSC genomes, 168 putative effectors were shared between all 18 genomes, and an additional 39 and 66 were specific for the *Fv* and *F. tenuicristatum* genomes, respectively (Fig. 6C). Most of the effectors in these groups were uncharacterized, the few characterized effectors of interest included multiple representatives of pectate lyase, glycosyl hydrolase, necrosis inducing protein, cellulose binding protein, beta-glucosidase, ribonuclease, esterase, and proteins with a LysM domain (Supplemental Table S12).

### Identification of differentially expressed genes and effectors in *Fv* 77-13-4 after pisatin treatment

Transcriptome analysis with Iso-seq (Wang et al. 2016) was used to identify genes differentially expressed in *Fv* 77-13-4 after treatment with the pea phytoalexin pisatin (Fig. 7). A principal component analysis (PCA) showed that the control DMSO and pisatin treated samples at T=0 clustered very close together, indicating minimal changes in the gene expression pattern between these sample conditions. However, the DMSO and pisatin treated samples collected 9 hours after treatment (T=9) clustered separately, indicating different patterns of gene expression (Fig. 7A). The differential gene expression analysis of the pisatin treated samples of *Fv* 77-13-4 revealed 96 genes (95 upregulated and 1 downregulated) differentially expressed at timepoint 0 hr (PT0) (Supplemental Table S13), and 1,155 genes (728 upregulated and 427 downregulated) differentially expressed at 9 hours after pisatin treatment (PT9) (Supplemental Table S14) when compared to the DMSO only treated samples at the corresponding timepoints (0hr (DT0) and 9hr (DT9)), respectively (Fig. 7B).

**Fig. 7:**
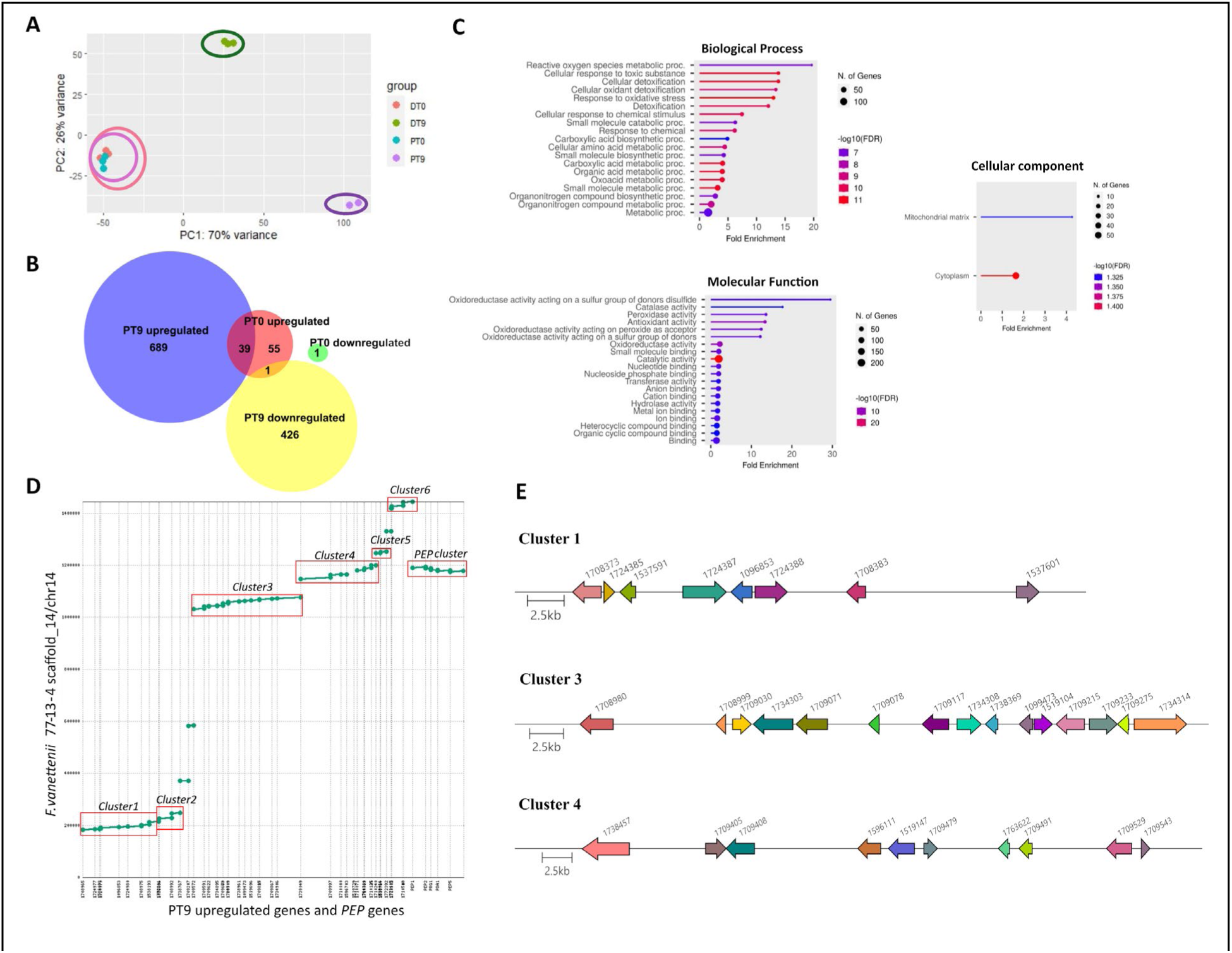
Differential gene expression in *F. vanettenii* 77-13-4 after pisatin treatment. **(A)** PCA plot displaying clustering as well as similarity between samples of various treatment conditions. **(B)** Venn diagram showing number of differentially expressed genes in PT0 and PT9 treatment conditions. **(C)** GO term analysis of upregulated genes at PT9. Top twenty significantly enriched GO terms at FDR < 0.05 for three GO categories: Biological Process, Molecular Function, and Cellular Component are shown if present **(D)** Dot plot showing MUMmer alignment of differentially upregulated genes at PT9 against *Fv* 77-13-4 chromosome 14. X-axis shows transcript Ids of upregulated genes at PT9. Potential gene clusters and the *PEP* cluster are highlighted with red rectangular boxes. **(E)** Plots showing organization of upregulated genes in top three large clusters. Each color represents different genes with arrows indicating their orientation in the scaffold. Protein Ids are indicated above each arrow. Figures were generated using the program Clinker.

In order to identify pathways and processes that were activated in *Fv* 77-13-4 after pisatin treatment, a gene enrichment analysis using ShinyGO (Ge et al. 2020) was performed for genes upregulated at nine hours after treatment. The GO term enrichment, KEGG pathway enrichment, as well as gene set enrichment analysis was performed for genes upregulated after pisatin treatment, where transcripts for genes involved in cellular detoxification and various metabolic processes (including reactive oxygen species) were significantly enriched (Fig. 7C and Supplemental Fig. S5). Similarly, transcripts for genes encoding oxidoreductase, catalase, and peroxidase activity, and binding to small molecules, nucleotides, and metal ions were significantly enriched (Fig. 7C and Supplemental Fig. S5). As for the cellular component, only transcripts for genes involved in the mitochondrial matrix and cytoplasm were significantly enriched (Fig. 7C). Overall, this gene enrichment analysis indicates that when the fungus is treated with pisatin, multiple pathways are activated that are directed at detoxifying the compound as well as other metabolic processes that enable it to survive in the presence of the phytoalexin.

Furthermore, analysis of the differentially expressed genes (DEGs) identified, of the 1,155 DEGs nine hours after treatment with pisatin, 38 were predicted to be effectors, including eight that were upregulated and 30 downregulated (Supplemental Fig. S6). The Pfam domain analysis of these 38 predicted effectors revealed that only ten proteins encoded functional domains including those targeting the plant cell wall, an upregulated putative glycosyl hydrolase 43 family member, and a pectate lyase (Supplemental Table S15). Interestingly, many of these differentially expressed effectors of *Fv* 77-13-4 were found to be clustered on core chromosomes, where six of the 38 resided on scaffold 6 (chromosome 6) and ten of the putative effectors resided on scaffold 7 (chromosome 7). The eight effectors upregulated after pisatin treatment were distributed across various scaffolds, whereas most of the downregulated effectors were distributed within close proximity on the same scaffolds (Supplemental Fig. S6).

Analysis of the genes upregulated at 9 hours after pisatin treatment revealed that of the 728 upregulated genes, 46 genes resided on chromosome 14 (scaffold_14). Interestingly, six gene clusters were upregulated on chromosome 14 with sizes ranging from ∼6 kb to 54 kb (Fig. 7D-E; Supplemental Table S16). The largest cluster (cluster 4, ∼54 kb) consisted of ten genes [six hypothetical proteins, two cytochrome P450s, one major facilitator transporter, and one Ubid decarboxylase] (Supplemental Table S16). In cluster 4, one of the cytochrome P450 enzymes (transcript ID 1519739) was predicted to possess a function related to pisatin demethylase, suggesting this could be another cluster on chromosome 14 related to virulence on pea (Supplemental Table S16).

## Discussion

Genomic features, virulence factors, secreted effector proteins, and accessory chromosomes of diverse fungal pathogens provide insight into their evolutionary relationships as well as host-specific characteristics. Long read genome sequencing and comparative analysis of sixteen isolates of *Fv* and five isolates of *Fop* collected from various host plants and geographic locations identified chromosomes and loci that could play an important role in causing disease on agriculturally important crops with an emphasis on garden pea.

Synteny analysis of the accessory genomes revealed that while there was diversity between the FSSC isolates, some syntenic regions were shared between the accessory genomes of 230-30-6, 77-13-4, T33, T77, T78, T161, T200, T219, and T299. When compared phylogenetically, with the exception of 230-30-6, these isolates clade together (Fig. 1A). Similarly, the accessory genomes of isolates T23 and T36 shared some syntenic regions and were also phylogenetically closely related. The presence of these syntenic regions within the accessory genomes and their close phylogenic relationship indicates that these regions have been vertically inherited. Despite the presence of the syntenic regions, there were multiple rearrangements indicating that the accessory genome of these fungi might be hotspots for recombination. Synteny analysis of the *PDA* containing scaffolds of the FSSC isolates identified four distinct groups with synteny. The difference in the phylogenetic relationship of 230-30-6 compared to other isolates within the syntenic group in accessory genome and *PDA* scaffolds may have resulted from multiple genetic crosses of this laboratory strain. Thus, the phylogeny may not accurately represent the original wildtype accessory genome of this strain. Moreover, since most of these isolates were acquired from different host plants and geographic locations, this data suggests that the accessory genome in *Fv* isolates might have been shaped by host-specific environmental factors that contributed to the diversity in these genetic regions.

Regions of the accessory genome were found to be shared between members of the FSSC and *Fop*, in particular scaffolds that were found to reside on the *PDA* encoding chromosome in *Fv* 77-13-4. This suggests that there is a shared origin for at least some of the accessory genome of *Fv* and *Fop* pea pathogens. One explanation for this similarity is the accessory genome was acquired in the last shared common ancestral fungus ∼65 Mya, and then vertically inherited in *Fv* and *Fop*. However, this would require the accessory genome to be lost in most, if not all, of the other Species Complexes of *Fusarium*. A more parsimonious explanation would be that each of these species inherited these accessory regions from either a common donor organism or were transferred between one another and have since diverged.

The presence / absence and synteny analysis of the *PEP* cluster revealed there is conservation of the *PEP* genes across FSSC isolates. Even though *PEP1* homologs were found in all FSSC genomes it was rarely found on the same *PDA* scaffold with other *PEP* genes. This suggests that while *PEP1* contributes to pathogenicity on pea (Han et al. 2001), it may not be a part of the *PEP* cluster for most of the genomes of FSSC isolates pathogenic on pea. This observation is in agreement with previous reports that *Fv* isolates can carry multiple homologs of *PEP1* (Temporini and VanEtten 2002; Coleman et al. 2009). The presence of multiple *PEP* genes in the genome of the pea pathogen *F. tenuicristatum* (NRRL 22470) suggests that the *PEP* cluster might have originated in this species through horizontal gene transfer (Temporini and VanEtten 2004).

Plant pathogens are required to overcome various host plant resistance mechanisms, one of them being the synthesis of antimicrobial compounds called phytoalexins (Jones and Dangl 2006; Ahuja et al. 2012). Garden pea in particular produces the phytoalexin pisatin in response to infection by various pathogens (Perrin and Bottomley 1961). The ability to detoxify pisatin by *Fv* is encoded by several alleles of the *PDA* genes (Maloney and VanEtten 1994), which encodes the cytochrome P450 Pda and detoxifies pisatin to the less toxic 6a-hydroxymaackiain and 3-hydroxymaackiain-isoflavan (Delserone et al. 1999). In the pisatin induced transcriptomic study of *Fv* 77-13-4, genes involved in cellular detoxification and/or cellular response to toxic substances are significantly upregulated in the pisatin treated samples. These significantly upregulated genes include different classes of transporters, cytochrome P450s, pectate lyase, glycosyl hydrolases, esterases, etc. The mechanisms of tolerance of various phytoalexins in fungi has been well documented and usually involve enzymatic detoxification and/or non-degradative tolerance (mediated by transporters) (VanEtten et al. 2001). In addition to the previously characterized *PEP* cluster, six other clusters of genes were found to be upregulated in response to pisatin treatment indicating there could be multiple tolerance mechanisms involved in pathogenicity. The pisatin treated transcriptome data clearly indicates that *Fv* is employing multiple tolerance mechanisms in response to pisatin.

The race of a fungal pathogen is defined as the biotype that has the same combination of virulence genes and is able to cause disease on a specific cultivar of a host species in a gene-for-gene interaction (Flor 1971). Four *Fop* races (1, 2, 5, and 6) have been described globally (Achari et al. 2021). Identification of effector proteins in *Fop* and comparative analysis based on their race enabled candidate effectors specific for each race to be identified that may serve as a race defining avirulence locus. These race specific effectors could be exploited to identify the corresponding resistance gene(s) in pea cultivars. Most of the *Fop* race specific effectors lacked functional domains, although the ones identified were a glycosyl hydrolase and lactonase as reported in a previous study (Achari et al. 2021). The presence of effectors encoding carbohydrate-active enzymes (CAZymes) such as glycosyl hydrolase, amidase, and beta-glucosidase suggests these proteins could be an important virulence strategy for particular races, facilitating host plant infection through degradation of plant polysaccharides. Similarly, some of these effectors encode a transcription factor domain that could be important for gene regulation as well as pathogen virulence. Having transcription factors specific to certain races could be an important host specificity strategy acquired by these particular races to regulate multiple genes as well as pathogen virulence.

As observed with the *Fop* effectors, conserved putative effectors in the FSSC genomes indicated cell wall degradation plays a key role in virulence. Putative cell wall degrading effectors included pectate lyase enzymes, cellulose binding proteins, glycosyl hydrolases, carbohydrate binding proteins, necrosis inducing proteins, esterase, and acyl hydrolase. Two pectate lyases (*PELA* and *PELD*) have previously been shown to be required for virulence on pea, although mutation of a single gene does not affect virulence (Rogers et al. 2000), indicating they are functionally redundant.

The genomes from *Fv* and *Fop* isolates provide insight into the genomic diversity and host specificity towards garden pea within two phylogenetically divergent species complexes. The diversity in the accessory genome across FSSC isolates illustrates the dynamic nature of these chromosomes, although implies a shared common origin. Shared accessory regions between the FSSC and *Fop* genomes were evident, indicating horizontal gene/chromosome transfer likely occurred in their evolutionary history. Analysis of the secreted proteins, pangenome, and TEs provides a better understanding of host-specific genomic features and lays a basis for broadening our knowledge on various pathogenicity factors within the FSSC and FOSC.

## Methods

### Selection of *F. vanettenii* and *F. oxysporum* f. sp. *pisi* isolates for this study

A diverse set of sixteen *Fv* isolates, five isolates of *Fop*, and *F. tenuicristatum* isolate NRRL 22470 collected from various geographic locations and host plants were used in this study (Table S1). These isolates were grown and maintained on V8 juice agar medium or minimum nutrient medium (M-100) (Stevens 1974). A previously assembled genome of the pea pathogen *Fop* HDV247 was also included in this study for comparative genomic analysis (Williams et al. 2016).

### DNA/RNA isolation, sequencing, genome assembly, and annotation

High quality genomic DNA was extracted from all *Fv* and *Fop* isolates using a CTAB method and a Qiagen genomic-tip (Kohler et al. 2011). RNA extraction was conducted as previously described (Coleman et al. 2009) except it was harvested from mycelia grown in two different media, M-100 and glucose-asparagine (GA) medium [20 g glucose, 3 g asparagine, 0.87 g K2HPO4, 0.14 g MgSO4.7H2O, and 1 mL Steinberg’s microelements diluted to 1 liter with H2O].

Genome sequencing, assembly, and annotation was performed for all isolates at the DOE JGI. All genomes except for *F. oxysporum* f. sp. *pisi* T415, *F. tenuicristatum* NRRL 22470, and *F. vanettenii* 230-30-6 were sequenced with PacBio, for which genomic DNA was sheared around 10 kbp (30-50 kbp for *F. vanettenii* 77-13-4) using the Megaruptor® 3 (Diagenode) or g-TUBE (Covaris). The sheared DNA was treated with exonuclease to remove single-stranded ends, DNA damage repair enzyme mix, end-repair/A-tailing mix and ligated with barcoded overhang adapters using SMRTbell Express Template Prep Kit 2.0 (PacBio) and purified with AMPure® PB Beads (PacBio) or Barcoded Adapter Plate 3.0 (PacBio) using SMRTbell Express Template Prep Kit 2.0 (PacBio). Libraries were size-selected using the 0.75% agarose gel cassettes with Marker S1 and High Pass protocol on the BluePippin (Sage Science). PacBio Sequencing primer was then annealed to the SMRTbell template library and sequencing polymerase was bound to them using Sequel II Binding kit. The prepared SMRTbell template libraries were then sequenced on a Pacific Biosystems’ Sequel II sequencer using sample dependent sequencing primer, 8M v1 SMRT cells, and Version 2.0 sequencing chemistry with 1×1800 sequencing movie run times. The artifact- and mitochondria-filtered CCS reads were assembled with Flye (Kolmogorov et al. 2019; https://github.com/fenderglass/Flye) either using - t 32 --pacbio-hifi and subsequently polished with two rounds of RACON version 1.4.13 racon with -u -t 36 (Vaser et al. 2017; https://github.com/lbcb-sci/racon), or, using -g 40M --asm-coverage 50 --pacbio-corr to generate an assembly and polished with gcpp --algorithm arrow version SMRTLINK v8.0.0.80529 (https://www.pacb.com/support/software-downloads). In all assemblies, contigs less than 1,000 bp were excluded.

The genomes of *F. oxysporum* f. sp. *pisi* T415, *F. tenuicristatum* NRRL 22470, and *F. vanettenii* 230-30-6 was sequenced using Illumina. For Illumina fragment libraries, 100 ng of DNA was sheared to 300 bp using the Covaris LE220 and size selected using SPRI beads (Beckman Coulter). The fragments were treated with end-repair, A-tailing, and ligation of Illumina compatible adapters (IDT, Inc) using the KAPA-Illumina library creation kit (KAPA biosystems). For Illumina long mate-pairs (LMP) libraries, 1.7ug of DNA was sheared using the Covaris g-TUBE(TM) and gel size selected for 7. The sheared DNA was treated with end repair and ligated with biotinylated adapters containing loxP. The adapter ligated DNA fragments were circularized via recombination by a Cre excision reaction (NEB). The circularized DNA templates were then randomly sheared using the Covaris LE220 (Covaris). The sheared fragments were treated with end repair and A-tailing using the KAPA-Illumina library creation kit (KAPA biosystems) followed by immobilization of mate pair fragments on strepavidin beads (Invitrogen). Illumina compatible adapters (IDT, Inc) were ligated to the mate pair fragments and 8 cycles of PCR was used to enrich for the final library (KAPA Biosystems). The prepared libraries were quantified using KAPA Biosystems’ next-generation sequencing library qPCR kit and run on a Roche LightCycler 480 real-time PCR instrument. Sequencing of the flowcell was performed on the Illumina NovaSeq sequencer using NovaSeq XP V1 reagent kits, S4 flowcell, following a 2×151 indexed run recipe.

Illumina reads filtered for artifact and process contamination and trimmed for quality were assembled with AllPathsLG version R49403 (Gnerre et al. 2011). For *F.tenuicristatum* NRRL 22470 with no LMP data, in silico LMP library with insert 3000 +/- 300 bp was produced from the initial assembly of fragment library reads with Velvet (Zerbino and Birney 2008) and then assembled with AllPathsLG.

For annotation, all transcriptomes were sequenced using Illumina RNA-Seq. Plate-based RNA sample prep was performed on the PerkinElmer Sciclone NGS robotic liquid handling system using Illumina’s TruSeq Stranded mRNA HT sample prep kit utilizing poly-A selection of mRNA following the protocol outlined by Illumina in their user guide: https://support.illumina.com/sequencing/sequencing_kits/truseq-stranded-mrna.html, and with the following conditions: total RNA starting material was 1000 ng per sample and 8 cycles of PCR was used for library amplification. The prepared libraries were quantified using KAPA Biosystems’ next-generation sequencing library qPCR kit and run on a Roche LightCycler 480 real-time PCR instrument. Sequencing of the flowcell was performed on the Illumina NovaSeq sequencer using NovaSeq XP V1 reagent kits, S4 flowcell, following a 2×151 indexed run recipe.

Using BBDuk (https://sourceforge.net/projects/bbmap/), raw RNA-Seq reads were evaluated for artifact sequence by kmer matching (kmer=25), allowing 1 mismatch and detected artifact was trimmed from the 3’ end of the reads. RNA spike-in reads, PhiX reads and reads containing any Ns were removed. Quality trimming was performed using the phred trimming method set at Q6. Finally, following trimming, reads under the length threshold were removed (minimum length 25 bases or 1/3 of the original read length - whichever is longer). Filtered reads were assembled into consensus sequences using Trinity v.2.11.0 (Grabherr et al. 2011).

Genome assemblies were annotated with transcriptome inputs using the JGI Fungal Annotation Pipeline (Grigoriev et al. 2014).

### Phylogenetic analysis of FSSC and *Fop* isolates

*TEF1*, *RPB1*, and *RPB2* gene sequences were identified using BLAST+ (Camacho et al. 2009) for all FSSC and *Fop* isolates. A multiple gene sequence alignment was conducted in MEGA11 (Tamura et al. 2021), and gaps and non-conserved regions were trimmed using Gblocks (Castresana 2000). Optimal model selection was done using PhyML (Lefort et al. 2017) and a maximum likelihood phylogeny tree with 1,000 bootstrap replicates was constructed using RAxML (Stamatakis 2014). Visualization of the tree was accomplished using iTOL (Letunic and Bork 2024).

### Prediction of effector proteins

The effector proteins in the FSSC and *Fop* genomes were predicted using a three-step pipeline. Initially, the secreted proteins were identified using SignalP 6.0 (Teufel et al. 2022), followed by filtering out proteins containing transmembrane domains using DeepTMHMM (Hallgren et al. 2022). From the list of the secreted proteins, the putative effector proteins were predicted using the program EffectorP 3.0 (Sperschneider and Dodds 2022). The final list of predicted effector proteins were classified into various orthogroups using the program OrthoFinder (Emms and Kelly 2019) and visualized via the R package UpSetR (Conway et al. 2017). Interesting orthogroups of these proteins were further analyzed to identify protein domains using the Pfam database (Mistry et al. 2021). The homologs of fourteen secreted in xylem (SIX) effectors and four early root colonization (ERC) effectors previously reported in *F. oxysporum* f. sp. *lycopersici* (Lievens et al. 2009; Redkar et al. 2022) were searched against the list of effectors identified in *Fop* genomes using BLAST+ (e-value <0.01; bitscore >50) (Camacho et al. 2009).

### RNA sequencing and differential gene expression analysis

The experimental setup and sample collection for the RNA sequencing study of *Fv* 77-13-4 was conducted as previously described (Liu et al., 2003) with a few modifications. Mycelium from *Fv* 77-13-4 was grown for 24 hours, collected by vacuum filtration and suspended in 0.05 M PBS buffer, pH 6.5. Pisatin dissolved in DMSO was added to the PBS-mycelia suspension at a final concentration of 31 μg/mL in 0.5% DMSO. Samples treated with only 0.5% DMSO were used as a negative control for each sample. Mycelium from individual treatments were collected at 0 hour and 9 hours after treatment and flash frozen at -80°C for RNA extraction. The RNA-seq experiment had 3 different biological replicates and each replicate had 4 sampling conditions (DMSO 0hr, Pisatin 0hr, DMSO 9hr and Pisatin 9hr indicated as DT0, DT9, PT0, and PT9, respectively). RNA extraction from the collected samples was done as described above. cDNA library preparation and RNA sequencing using the PacBio Iso-seq method was performed at the JGI, as was processing the RNAseq reads and de novo transcriptome assembly. The PCA analysis revealed that the pisatin treated sample after 9 hours from replicate 1 (R1PT9) clustered separately and was an outlier for the group and therefore was removed from further analysis. The differential gene expression analysis was performed by comparing the pisatin treated samples with DMSO only treated samples for each timepoint using the Bioconductor package DESeq2 (Love et al., 2014). The genes with a log fold change threshold > 1 and p.adj <0.05 were considered as differentially expressed. The gene enrichment analysis of the differentially expressed genes was performed using the program ShinyGO 0.80 (Ge et al. 2020) where the GO enrichment pathways, KEGG enrichment pathways, and gene set enrichment analysis (GSEA) was performed and visualized. The pathways and gene sets with a false discovery rate (FDR) correction at p < 0.05 were considered as significantly enriched. The differentially upregulated genes at 9hr were further mapped against chromosome 14 using MUMmer 3.22 (Kurtz et al. 2004) to identify the potential gene clusters upregulated after pisatin treatment.

The list of genes that were differentially expressed at both 0hr and 9hr were further analyzed to identify secreted effector proteins using the pipeline as described above and the protein family domains were identified using the Pfam database (Mistry et al. 2021).

### Defining core and accessory genomic regions of FSSC and *Fop* isolates

The previously characterized genome assembly of *Fv* 77-13-4 (Coleman et al. 2009) was used as a reference to identify core and accessory (or lineage-specific) regions in the FSSC genomes using MUMmer 3.22 (Kurtz et al. 2004). Similarly the *F. oxysporum* f. sp. *vasinfectum* (*Fov*) TF1 genome assembly (Seo et al. 2020) was used as a reference to identify core and accessory regions in *Fop* isolates. All FSSC and *Fop* genomes were aligned to the core genome of *Fv* 77-13-4 and *Fov* TF1, respectively, using MUMmer 3.22 (PROmer -mum, delta-filter -g - i 90) (Kurtz et al. 2004; Williams et al. 2016). The output of MUMmer was written into a bed file and the percentage of coverage of query scaffolds to reference scaffolds was calculated using Bedtools (genomecov) (Quinlan and Hall 2010). The scaffolds in the query genomes with ≥30% coverage were considered as core and the remaining scaffolds were considered as accessory (Williams et al. 2016). Since the genome of *F. tenuicristatum* strain NRRL 22470 was highly fragmented, the cutoffs for the MUMmer alignment were 70% identity (delta-filter -i 70) and genome coverage cutoff was ≥20%. In order to identify and visualize the conservation of accessory genome across FSSC and *Fop* isolates, the mummer2circos package was used (https://github.com/metagenlab/mummer2circos). In order to identify large syntenic blocks across the accessory scaffolds in the FSSC genomes, the FindSynteny function (minScore = 100000, maxGap = 10000, maxsep = 10000) in the DECIPHER R package (Wright 2016) was used.

### GO enrichment analysis of core and accessory proteins

The pangenome of all *Fop* and FSSC isolates used in this study was evaluated using the core and accessory proteins. The InterProScan functional annotation of all proteins from the genomes were obtained from JGI and separated into core and accessory. ShinyGO 0.80 (Ge et al. 2020) was used to perform an enrichment analysis of all core and accessory proteins. The proteins from FSSC isolates were compared against the *Fv* 77-13-4 v2 genome (Coleman et al. 2009) and the proteins from *Fop* isolates were compared against *F*. *oxysporum* f. sp. *lycopersici* 4287 (Ma et al. 2010; Ayhan et al. 2018) available in the ShinyGO database. The GO terms with FDR correction at p < 0.05 were considered as significantly enriched.

### Identification of *PEP* genes and *PDA* scaffolds in FSSC and *Fop* isolates

The *PEP* genes in all the FSSC and *Fop* genomes were identified using BLAST+ (Camacho et al. 2009) by generating a custom database of *PEP* gene sequences from *Fv* 77-13- 4. The scaffold having a significant hit to the *PDA1* gene was considered as a *PDA* scaffold. Microsynteny analysis of the *PEP* genes on the *PDA* scaffolds was conducted using the program SimpleSynteny (Veltri et al. 2016). Identification of syntenic blocks across the FSSC *PDA* scaffolds was performed using the FindSynteny function in the DECIPHER R package (Wright 2016).

### Identification of transposable elements in *Fop* and FSSC isolates

The *de novo* annotation of transposable elements (TEs) was done in all *Fop* and FSSC isolates using EDTA with the higher sensitivity setting (--sensitive 1) (Ou et al. 2019). The prediction of TEs by the Extensive *de novo* TE Annotator (EDTA) program (Ou et al. 2019), which is based on structural (to identify intact TEs) and coding (to classify them to families) features. TEs that lacked homology and coding features were labelled as ‘unknown’ TEs. The output from EDTA was aggregated, manually curated, and visualized using the stacked histogram.

## Data access

The data described in the publication can be accessed on the JGI Genome Portal and the JGI MycoCosm (Grigoriev et al. 2014). Data generated in this study has been submitted to the NCBI Bioproject database (https://www.ncbi.nlm.nih.gov/bioproject) under accession numbers PRJNA334374, PRJNA334357, PRJNA655082, PRJNA710761, PRJNA1080754, PRJNA1080755, PRJNA1080756, PRJNA1080757, PRJNA1080758, PRJNA1080773, PRJNA1080774, PRJNA1080800, PRJNA1080801, PRJNA1080802, PRJNA1080803, PRJNA1080804, PRJNA1080805, PRJNA1080809, PRJNA1080810, PRJNA1080811, PRJNA1080812, PRJNA1080813, PRJNA1080814, and PRJNA1080878, for genome sequence and PRJNA655083, PRJNA572453, PRJNA538191, PRJNA710675, PRJNA1235803, PRJNA710674, PRJNA1235819, PRJNA1235836, PRJNA1235833, PRJNA1235837, PRJNA1235831, PRJNA1232534, PRJNA1235827, PRJNA1235806, PRJNA1235826, PRJNA1235801, PRJNA1235828, PRJNA1235825, PRJNA1235802, PRJNA1235811, PRJNA1235830, PRJNA1235817, PRJNA1235820, for transcriptome data used to aid in annotation, and PRJNA1235816, PRJNA1235814, PRJNA1235821, PRJNA1235808, PRJNA1235812, PRJNA1235813, PRJNA1235815, PRJNA1235804, PRJNA1235797, PRJNA1235805, PRJNA1235807, and PRJNA1235798 for Iso-seq.

## Acknowledgments

Genome sequencing and assembly (proposals 10.46936/10.25585/6001402 and 10.46936/10.25585/60001201) was conducted by the U.S. Department of Energy (DOE) Joint Genome Institute (https://ror.org/04xm1d337), a DOE Office of Science User Facility, and is supported by the Office of Science of the DOE under contract number DE-AC02-05CH11231. A.P. and J.J.C. were supported by the USDA Hatch Program.

## Supplemental Figure legends

**Fig. S1: Comparison of basic genomic features between core and accessory genomic regions of *Fop* and FSSC isolates. (A)** Genome size (Mbp), GC content (%) and number of genes for *Fop* isolates. **(B)** Genome size (Mbp), GC content (%), and number of genes respectively for FSSC isolates.

**Fig. S2: Gene enrichment analysis of core and accessory genomic regions of *Fop* isolates.** Top ten significantly enriched gene ontology pathways of the core and accessory genome in *Fop* isolates. The statistical significance of gene enrichment was determined at FDR threshold < 0.05. The enriched pathways are categorized as biological process (BP), molecular function (MF), and cellular components (CC). *None of the gene ontology pathways under cellular component category were significantly enriched in the accessory genome of *Fop* isolates.

**Fig. S3: Synteny analysis of *PDA* scaffolds in FSSC genomes (A)** Synteny dotplot from pairwise comparison of *PDA* scaffolds of all FSSC isolates. **(B)** – **(E)** Synteny dotplots of pairwise comparison of *PDA* scaffolds from a subset of FSSC isolates. Plots above diagonal with black lines indicate syntenic regions in the same orientation and red lines indicate inversions. Plots below diagonal indicate synteny based on score, green (highest) to blue to magenta (lowest). Synteny dotplots were created using DECIPHER package in R. All possible syntenic blocks > 10 kb are shown.

**Fig. S4: Synteny analysis of accessory genomes between the FSSC (A)** Synteny dotplot from pairwise comparison of accessory genome in all FSSC isolates. **(B)** – **(C)** Synteny dotplots of pairwise comparison of accessory genome from a subset of FSSC isolates. Plots above diagonal with black lines indicate syntenic regions in same orientation and red lines indicate inversions. Plots below diagonal indicate synteny based on score, green (highest) to blue to magenta (lowest). Synteny dotplots were created using DECIPHER package in R. Only syntenic blocks > 100 kb are shown.

**Fig. S5: KEGG and gene set enrichment analysis (GSEA) of genes upregulated after pisatin treatment. (A)** Top twenty significantly enriched KEGG pathways of upregulated genes at PT9 treatment conditions with FDR < 0.05. **(B)** Gene set enrichment analysis of genes upregulated at PT9 treatment condition. Only top 20 gene sets are shown that are significant at FDR < 0.05.

**Fig. S6: Prediction of differentially expressed effector proteins after pisatin treatment. (A)** Venn diagram showing distribution of effector proteins in various FSSC groups and effectors differentially expressed 9 hours after pisatin treatment. **(B)** Dot plot showing the MUMmer alignment of differentially expressed effectors in 9-hour pisatin treatment condition against *Fv* 77-13-4 genome. Red circles indicate the upregulated effectors after treatment with pisatin.

## References

Achari SR, Edwards J, Mann RC, Kaur JK, Sawbridge T, Summerell BA. 2021. Comparative transcriptomic analysis of races 1, 2, 5 and 6 of *Fusarium oxysporum* f.sp. *pisi* in a susceptible pea host identifies differential pathogenicity profiles. BMC Genom 22: 734.

Ahuja I, Kissen R, Bones AM. 2012. Phytoalexins in defense against pathogens. Trends Plant Sci 17: 73–90.

Ayhan DH, López-Díaz C, Di Pietro A, Ma L-J. 2018. Improved assembly of reference genome *Fusarium oxysporum* f. sp. *lycopersici* strain fol4287. Microbiology Resource Announcements 7: 10.1128/mra.00910-18.

Camacho C, Coulouris G, Avagyan V, Ma N, Papadopoulos J, Bealer K, Madden TL. 2009. BLAST+: architecture and applications. BMC Bioinform 10: 421.

Castresana J. 2000. Selection of conserved blocks from multiple alignments for their use in phylogenetic analysis. Mol Biol Evol 17: 540–552.

Coleman JJ. 2016. The *Fusarium solani* species complex: ubiquitous pathogens of agricultural importance. Mol Plant Pathol 17: 146–158.

Coleman JJ, Rounsley SD, Rodriguez-Carres M, Kuo A, Wasmann CC, Grimwood J, Schmutz J, Taga M, White GJ, Zhou S, et al. 2009. The genome of *Nectria haematococca*: Contribution of supernumerary chromosomes to gene expansion. PLoS Genet 5: e1000618.

Coleman JJ, Wasmann CC, Usami T, White GJ, Temporini ED, McCluskey K, VanEtten HD. 2011. Characterization of the gene encoding pisatin demethylase (*FoPDA1*) in *Fusarium oxysporum*. MPMI 24: 1482–1491.

Conway JR, Lex A, Gehlenborg N. 2017. UpSetR: an R package for the visualization of intersecting sets and their properties. Bioinformatics 33: 2938–2940.

Delserone LM, McCLUSKEY K, Matthews DE, Vanetten HD. 1999. Pisatin demethylation by fungal pathogens and nonpathogens of pea: association with pisatin tolerance and virulence. Physiol Mol Plant Pathol 55: 317–326.

Emms DM, Kelly S. 2019. OrthoFinder: phylogenetic orthology inference for comparative genomics. Genome Biol 20: 238.

Flor HH. 1971. Current status of the gene-for-gene concept. Annual Review of Phytopathology 9: 275–296.

Funnell DL, Matthews PS, VanEtten HD. 2001. Breeding for highly fertile isolates of *Nectria haematococca* MPVI that are highly virulent on pea and in planta selection for virulent recombinants. Phytopathology® 91: 92–101.

Ge SX, Jung D, Yao R. 2020. ShinyGO: a graphical gene-set enrichment tool for animals and plants. Bioinformatics 36: 2628–2629.

Geiser DM, Al-Hatmi AMS, Aoki T, Arie T, Balmas V, Barnes I, Bergstrom GC, Bhattacharyya MK, Blomquist CL, Bowden RL, et al. 2021. Phylogenomic analysis of a 55.1-kb 19-gene dataset resolves a monophyletic *Fusarium* that includes the *Fusarium solani* species complex. Phytopathology 111: 1064–1079.

Gnerre S, MacCallum I, Przybylski D, Ribeiro FJ, Burton JN, Walker BJ, Sharpe T, Hall G, Shea TP, Sykes S, et al. 2011. High-quality draft assemblies of mammalian genomes from massively parallel sequence data. Proc Natl Acad Sci U S A 108: 1513–1518.

Grabherr MG, Haas BJ, Yassour M, Levin JZ, Thompson DA, Amit I, Adiconis X, Fan L, Raychowdhury R, Zeng Q, et al. 2011. Trinity: reconstructing a full-length transcriptome without a genome from RNA-Seq data. Nat Biotechnol 29: 644–652.

Grigoriev IV, Nikitin R, Haridas S, Kuo A, Ohm R, Otillar R, Riley R, Salamov A, Zhao X, Korzeniewski F, et al. 2014. MycoCosm portal: gearing up for 1000 fungal genomes. Nucleic Acids Research 42: D699–D704.

Hallgren J, Tsirigos KD, Pedersen MD, Armenteros JJA, Marcatili P, Nielsen H, Krogh A, Winther O. 2022. DeepTMHMM predicts alpha and beta transmembrane proteins using deep neural networks. 2022.04.08.487609. https://www.biorxiv.org/content/10.1101/2022.04.08.487609v1 (Accessed February 22, 2024).

Han YN, Liu XG, Benny U, Kistler HC, VanEtten HD. 2001. Genes determining pathogenicity to pea are clustered on a supernumerary chromosome in the fungal plant pathogen *Nectria haematococca*. Plant J 25: 305–314.

Jones JDG, Dangl JL. 2006. The plant immune system. Nature 444: 323–329.

Kohler A, Murat C, Costa M. 2011. Extraction of high quality DNA for genome sequencing. INRA Nancy Equipe Ecogénomique, Champenoux, France 4.

Kolmogorov M, Yuan J, Lin Y, Pevzner PA. 2019. Assembly of long, error-prone reads using repeat graphs. Nat Biotechnol 37: 540–546.

Kurtz S, Phillippy A, Delcher AL, Smoot M, Shumway M, Antonescu C, Salzberg SL. 2004. Versatile and open software for comparing large genomes. Genome Biol 5: R12.

Lefort V, Longueville J-E, Gascuel O. 2017. SMS: smart model selection in PhyML. Mol Biol Evol 34: 2422–2424.

Letunic I, Bork P. 2024. Interactive Tree of Life (iTOL) v6: recent updates to the phylogenetic tree display and annotation tool. Nucleic Acids Res 52: W78–W82.

Lievens B, Houterman PM, Rep M. 2009. Effector gene screening allows unambiguous identification of *Fusarium oxysporum* f. sp. *lycopersici* races and discrimination from other formae speciales. FEMS Microbiology Letters 300: 201–215.

López Díaz C, Ayhan DH, Rodríguez López A, Gómez Gil L, Ma L-J, Di Pietro A. 2025. Transposons and accessory genes drive adaptation in a clonally evolving fungal pathogen. Nat Commun 16: 6982.

Ma L-J, van der Does HC, Borkovich KA, Coleman JJ, Daboussi M-J, Di Pietro A, Dufresne M, Freitag M, Grabherr M, Henrissat B, et al. 2010. Comparative genomics reveals mobile pathogenicity chromosomes in *Fusarium*. Nature 464: 367–373.

Maloney AP, VanEtten HD. 1994. A gene from the fungal plant pathogen *Nectria haematococca* that encodes the phytoalexin-detoxifying enzyme pisatin demethylase defines a new cytochrome P450 family. Mol Gen Genet 243: 506–514.

Miao VP, Covert SF, VanEtten HD. 1991. A fungal gene for antibiotic-resistance on a dispensable (“B”) chromosome. Science 254: 1773–1776.

Michielse CB, Rep M. 2009. Pathogen profile update: *Fusarium oxysporum*. Molecular Plant Pathology 10: 311–324.

Mistry J, Chuguransky S, Williams L, Qureshi M, Salazar GA, Sonnhammer ELL, Tosatto SCE, Paladin L, Raj S, Richardson LJ, et al. 2021. Pfam: the protein families database in 2021. Nucleic Acids Res 49: D412–D419.

Navasca A, Singh J, Rivera-Varas V, Gill U, Secor G, Baldwin T. 2025. Dispensable genome and segmental duplications drive the genome plasticity in *Fusarium solani*. Front Fungal Biol 6: 1432339.

O’Donnell K, Rooney AP, Proctor RH, Brown DW, McCormick SP, Ward TJ, Frandsen RJN, Lysøe E, Rehner SA, Aoki T, et al. 2013. Phylogenetic analyses of *RPB1* and *RPB2* support a middle Cretaceous origin for a clade comprising all agriculturally and medically important fusaria. Fungal Genetics and Biology 52: 20–31.

Ou S, Su W, Liao Y, Chougule K, Agda JRA, Hellinga AJ, Lugo CSB, Elliott TA, Ware D, Peterson T, et al. 2019. Benchmarking transposable element annotation methods for creation of a streamlined, comprehensive pipeline. Genome Biology 20: 275.

Perrin DR, Bottomley W. 1961. Pisatin: an antifungal substance from *Pisum sativum* L. Nature 191: 76–77.

Pokhrel A, Coleman JJ. 2024. Transcriptional enhancement of pathogenicity genes in the *PEP* cluster of *Fusarium vanettenii* is influenced by a pisatin-responsive gene (*PRG1*) and contributes to virulence on garden pea. Physiological and Molecular Plant Pathology 134: 102411.

Quinlan AR, Hall IM. 2010. BEDTools: a flexible suite of utilities for comparing genomic features. Bioinformatics 26: 841–842.

Redkar A, Sabale M, Schudoma C, Zechmann B, Gupta YK, López-Berges MS, Venturini G, Gimenez-Ibanez S, Turrà D, Solano R, et al. 2022. Conserved secreted effectors contribute to endophytic growth and multihost plant compatibility in a vascular wilt fungus. Plant Cell 34: 3214–3232.

Rodriguez-Carres A, White G, Tsuchiya D, Taga M, VanEtten HD. 2008. The supernumerary chromosome of *Nectria haematococca* that carries pea-pathogenicity-related genes also carries a trait for pea rhizosphere competitiveness. Appl Environ Microbiol 74: 3849–3856.

Rogers LM, Kim Y-K, Guo W, González-Candelas L, Li D, Kolattukudy PE. 2000. Requirement for either a host- or pectin-induced pectate lyase for infection of *Pisum sativum* by *Nectria hematococca*. PNAS 97: 9813–9818.

Schmidt SM, Houterman PM, Schreiver I, Ma L, Amyotte S, Chellappan B, Boeren S, Takken FLW, Rep M. 2013. MITEs in the promoters of effector genes allow prediction of novel virulence genes in *Fusarium oxysporum*. BMC Genom 14: 119.

Seo S, Pokhrel A, Coleman JJ. 2020. The genome sequence of five genotypes of *Fusarium oxysporum* f. sp. *vasinfectum*: a resource for studies on *Fusarium* wilt of cotton. Mol Plant-Microbe Interact 33: 138–140.

Sperschneider J, Dodds PN. 2022. EffectorP 3.0: prediction of apoplastic and cytoplasmic effectors in fungi and oomycetes. MPMI 35: 146–156.

Stamatakis A. 2014. RAxML version 8: a tool for phylogenetic analysis and post-analysis of large phylogenies. Bioinformatics 30: 1312–1313.

Stevens RB, ed. 1974. Mycology guidebook. University of Washington Press, Seattle.

Tamura K, Stecher G, Kumar S. 2021. MEGA11: molecular evolutionary genetics analysis version 11. Mol Biol Evol 38: 3022–3027.

Temporini ED, VanEtten HD. 2004. An analysis of the phylogenetic distribution of the pea pathogenicity genes of *Nectria haematococca* MPVI supports the hypothesis of their origin by horizontal transfer and uncovers a potentially new pathogen of garden pea: *Neocosmospora boniensis*. Curr Genet 46: 29–36.

Temporini ED, VanEtten HD. 2002. Distribution of the pea pathogenicity (*PEP*) genes in the fungus *Nectria haematococca* mating population VI. Curr Genet 41: 107–114.

Teufel F, Almagro Armenteros JJ, Johansen AR, Gíslason MH, Pihl SI, Tsirigos KD, Winther O, Brunak S, von Heijne G, Nielsen H. 2022. SignalP 6.0 predicts all five types of signal peptides using protein language models. Nat Biotechnol 40: 1023–1025.

Van Dam P, de Sain M, Ter Horst A, van der Gragt M, Rep M. 2018. Use of comparative genomics-based markers for discrimination of host specificity in *Fusarium oxysporum*. Appl Environ Microbiol 84: e01868–17.

VanEtten H, Temporini E, Wasmann C. 2001. Phytoalexin (and phytoanticipin) tolerance as a virulence trait: why is it not required by all pathogens? Physiol Mol Plant Pathol 59: 83–93.

Vaser R, Sović I, Nagarajan N, Šikić M. 2017. Fast and accurate de novo genome assembly from long uncorrected reads. Genome Res 27: 737–746.

Veltri D, Wight MM, Crouch JA. 2016. SimpleSynteny: a web-based tool for visualization of microsynteny across multiple species. Nucleic Acids Res 44: W41–W45.

Wang B, Tseng E, Regulski M, Clark TA, Hon T, Jiao Y, Lu Z, Olson A, Stein JC, Ware D. 2016. Unveiling the complexity of the maize transcriptome by single-molecule long-read sequencing. Nat Commun 7: 11708.

Williams AH, Sharma M, Thatcher LF, Azam S, Hane JK, Sperschneider J, Kidd BN, Anderson JP, Ghosh R, Garg G, et al. 2016. Comparative genomics and prediction of conditionally dispensable sequences in legume-infecting *Fusarium oxysporum formae speciales* facilitates identification of candidate effectors. BMC Genom 17: 191.

Wright E S. 2016. Using DECIPHER v2.0 to analyze big biological sequence data in R. The R Journal 8: 352.

Yang H, Yu H, Ma L-J. 2020. Accessory chromosomes in *Fusarium oxysporum*. Phytopathology 110: 1488–1496.

Zerbino DR, Birney E. 2008. Velvet: Algorithms for de novo short read assembly using de Bruijn graphs. Genome Res 18: 821–829.

